# Voluntary running does not increase capillary blood flow but promotes neurogenesis and short-term memory in the APP/PS1 mouse model of Alzheimer’s disease

**DOI:** 10.1101/2020.02.28.969840

**Authors:** Kaja Falkenhain, Nancy E. Ruiz-Uribe, Mohammad Haft-Javaherian, Muhammad Ali, Stall Catchers contributors, Pietro E. Michelucci, Chris B. Schaffer, Oliver Bracko

## Abstract

Exercise exerts a beneficial effect on the major pathological and clinical symptoms associated with Alzheimer’ s disease in humans and mouse models of the disease. While numerous mechanisms for such benefits from exercise have been proposed, a clear understanding of the causal links remains elusive. Recent studies also suggest that cerebral blood flow in the brain of both Alzheimer’ s patients and mouse models of the disease is decreased and that the cognitive symptoms can be improved when blood flow is restored. We therefore hypothesized that the mitigating effect of exercise on the development and progression of Alzheimer’ s disease may be mediated through an increase in the otherwise reduced brain blood flow. To test this idea, we examined the impact of three months of voluntary wheel running in ∼1-year-old APP/PS1 mice on short-term memory function, brain inflammation, amyloid deposition, and cerebral blood flow. Our findings that exercise led to improved memory function, a trend toward reduced brain inflammation, markedly increased neurogenesis in the dentate gyrus, and no changes in amyloid-beta deposits are consistent with other reports on the impact of exercise on the progression of Alzheimer’ s related symptoms in mouse models. Notably, we did not observe any impact of wheel running on overall cortical blood flow nor on the incidence of non-flowing capillaries, a mechanism we recently identified as one contributing factor to cerebral blood flow deficits in mouse models of Alzheimer’ s disease. Overall, our results replicate previous findings that exercise is able to ameliorate certain aspects of Alzheimer’ s disease pathology, but show that this benefit does not appear to act through increases in cerebral blood flow.

## INTRODUCTION

### BACKGROUND

Alzheimer’ s disease (AD) is the most common neurodegenerative disease, showing increasing prevalence with age among the elderly population. In addition to a progressive decline in cognitive ability and memory, the disease is characterized by pathological changes that include the deposition of the protein fragment amyloid-beta (Aβ) to form extracellular amyloid plaques, the aggregation of hyperphosphorylated tau protein in neurofibrillary tangles, increased brain inflammation, reduced synaptic plasticity, and neuronal cell death [1]. Whereas less than 1% of AD cases are attributable to genetic mutations in the genes coding for amyloid precursor protein (APP), presenilin-1 (PS1), or presenilin-2 (PS2) that contribute to the early onset form of the disease, the vast majority of AD cases are sporadic, and have a later disease onset. Despite tremendous efforts, there are currently no effective treatments available.

### EFFECTS OF EXERCISE IN ALZHEIMER’ S DISEASE

Regular physical activity is recognized as providing health-enhancing benefits, including through reducing the incidence and severity of dementia [2-5]. Aerobic exercise interventions were shown to maintain and improve neurocognitive function in ageing individuals, particularly for executive-control function [6-9]. In multiple transgenic mouse models of Alzheimer’ s disease, exercise has been shown to affect AD pathology through multiple pathways. In particular, exercise in the form of wheel running was shown to lead to cognitive improvements, such as enhanced memory function, a rescue of synaptic plasticity, and an amelioration of some pathological features, including a decrease in levels of Aβ in some studies [10-17]. The latter was proposed to be mediated through changes in APP processing [18] and an increase in clearance of Aβ [19], possibly resulting from an elevation of Aβ clearance proteins [20] and increased glymphatic clearance [21]. In addition, expression of other genes linked to AD progression such as β-site APP-cleaving enzyme 1 (BACE1), presenilin 1 (PS1), insulin-degrading enzyme (IDE), tau, and the receptor for advanced glycation end products (RAGE) are reduced as a consequence of exercise [22, 23]. This may be due to a decrease in lipid-raft formation [23], activation of SIRT1 (sirtuin (silent mating type information regulation 2 homolog) 1) [12, 24], and an increase in autophagy [19, 25]. In addition, exercise has been proposed to mitigate mitochondrial dysfunction [11, 26], delay disease-related white matter volume loss [27-30], reduce neuroinflammation [19, 21, 22, 31-34] and oxidative stress [10, 19, 35], repress neuronal cell death [36], and favor adult neurogenesis [17, 37-40].

### BRAIN BLOOD FLOW IN ALZHEIMER’ S DISEASE

Cerebrovascular dysfunction has been described as an early feature in the development of many neurodegenerative diseases, including Alzheimer’ s disease [41]. For example, cerebral blood flow (CBF) was shown to be decreased by ∼30% in both patients with Alzheimer’ s disease and mouse models of APP overexpression [42, 43]. Other structural and functional cerebrovascular changes that have been described in AD include diminution in blood vessel diameter [44], loss of microvasculature [45], and impaired blood flow increases in response to neuronal stimuli or changes in blood pressure [46, 47]. Cardiovascular risk factors, such as hypertension, type 2 diabetes, and obesity [41], are also among the strongest predictors of AD risk and severity, again suggesting a tight association between vascular health and dementia.

Reduced cerebral blood flow could, in turn, exacerbate aspects of AD progression and independently contribute to cognitive impairment through, for example, acceleration of Aβ aggregation and plaque growth [48], causing white matter deficits in a tau mouse model [44], attenuating interstitial fluid flow [48], and a general cellular adaption to dysregulated blood flow, including pericytes dysfunction [47], metabolic rate contributions [49], and causing cortical atrophy [44]. Even though a clear understanding of the underlying causes and consequences of CBF reduction in Alzheimer’ s disease has not yet emerged, proposed mechanisms for this hypoperfusion include degeneration of pericytes [47, 50-52], a decrease in nitric oxide [53], Aβ-mediated constriction of capillaries by pericytes [54], and hypercoagulability due to fibrin interactions with Aβ [55, 56]. Additionally, a recent study identified an increased number of cortical capillaries showing obstructed blood flow due to neutrophil adhesion in capillary segments as a novel cellular mechanism underlying the reduction in cortical blood flow [57]. Interestingly, preventing adhesion of neutrophils rapidly increased CBF and led to improved memory performance within hours [57]. An acute improvement in memory performance due to CBF increase was seen in APP/PS1 mice as old as 16 months [58], suggesting that increasing brain blood flow can contribute to improved cognitive performance even in late stages of disease progression.

Exercise has been shown to cause a number of beneficial changes in blood vessels, including increased baseline velocity [59], enhanced cerebral vasodilator responses [60], reduced vascular oxidative stress [61], increased vascular density [62], and improved arterial pressure [63]. Therefore, we hypothesized that the mitigating effect of exercise observed in Alzheimer’ s disease may be mediated through an increase in the otherwise reduced brain blood flow. In particular, we sought to investigate the effects of three months of voluntary wheel running on CBF, memory function, and brain pathology in ∼1-year-old APP/PS1 mice.

## METHODS

### ANIMALS

All animal procedures were approved by the Cornell Institution Animal Care and Use Committee and were performed under the guidance of the Cornell Center for Animal Resources and Education (CARE). We used APP/PS1 transgenic mice (B6.Cg-Tg (APPswe, PSEN1dE9) 85Dbo/J; MMRRC_034832-JAX, The Jackson Laboratory) as mouse models of AD. This double transgenic mouse model expresses a chimeric mouse/human amyloid precursor protein (Mo/HuAPP695swe) and contains a mutant human transgene for presenilin 1 (PSE-dE9). Animals were of both sexes and ranged from 9 to 14 months in age at the beginning of the study.

### EXERCISE PARADIGM AND MONITORING

Mice were randomly assigned to either standard housing (sedentary group) or to a cage equipped with a low-profile wireless running wheel (ENV-047; Med Associates, Inc.) to which they had unrestricted access (running group), for a period of three months. A Wireless USB Hub (DIG-807; Med Associates, Inc.) and Wheel Manager Software (SOF-860; Med Associates, Inc.) were used to record running data from the wheels. All mice were single-housed and given free access to water and food.

### BEHAVIOR EXPERIMENTS

All experiments were performed under red lighting in an isolated room. Animals were taken into the procedure room for 1 h on each of two days preceding the experiment and were allowed to acclimate for 1 h prior to behavior sessions each day. The position of the mouse’ s head, body, and tail was automatically tracked by Viewer III software (Biobserve). In addition to the automatic results obtained by the software, a blinded researcher independently scored the mouse behavior manually. The testing arena was cleaned with 70% ethanol prior to use and in between each of the trials to remove any scent clues.

### OPEN FIELD TEST

The open field (OF) test was used to evaluate anxiety-like behavior and general locomotor ability. Mice were placed in the arena individually and allowed to freely explore for a single 30 min period during which time the tracking software recorded movement. Total track length, average velocity, and time spent in the inner and outer zones (defined as within 5 cm of the wall of the arena) of the arena were quantified. At the end of the test period, the mice were returned to their home cage.

### OBJECT REPLACEMENT TEST

The object replacement (OR) test was used to measure spatial recognition memory performance. Mice were presented with two identical objects and allowed to freely explore the objects for 10 min in a testing arena in which each wall was marked with a different pattern. Subsequently, mice were returned to their home cage for 60 min. Mice were then returned to the arena for a 5 min testing period with one of the objects moved to a different location. The object was moved such that both the intrinsic relationship between the objects and the position relative to patterned walls were altered. Exploration time for the objects was defined as any time when there was physical contact with an object or when the animal was oriented toward the object at a distance of 2 cm or less. Climbing over an object was not considered to be an explorative behavior unless that action was accompanied by directing the nose toward that object. The preference score was determined from the testing period data and was calculated as (exploration time of the replaced object/exploration time of both objects).

### Y-MAZE SPONTANEOUS ALTERNATION TEST

The Y-maze task was used to evaluate spatial working memory and willingness to explore new environments. Testing occurred in a Y-shaped maze consisting of three arms at 120° and made of light gray plastic. Each arm was 6 cm wide and 36 cm long, and had 12.5 cm high walls. Mice were placed in the Y-maze individually and allowed to freely explore the three arms for 6 min, during which time the tracking software recorded the mouse’ s movement. A mouse was considered to have entered an arm if all four limbs entered it, and to have excited it if all four limbs exited the arm. The number of arm entries and the number of consecutive entries into three different arms (alternating triad) was recorded to quantify the percentage of spontaneous alternation. Because the maximum number of alternating triads is equal to the total number of arm entries minus 2, the spontaneous alternation score was calculated as (number of alternating triads/[total number of arm entries −2]).

### NOVEL OBJECT RECOGNITION TEST

The novel object recognition test (NOR) was used to evaluate object-identity memory and explorative behavior. The testing protocol was identical to the object replacement test, except mice were returned to their home cage for 90 min in between trials and one of the initial objects was replaced, at the same location, with a novel object for the testing period. The preference score was analogously determined from the testing period data and calculated was as (exploration time of the novel object/exploration time of both objects).

### SURGICAL PREPARATION

For cranial window implantation, mice were anesthetized under 3% isoflurane and then maintained at 1.5 – 2% isoflurane in 100% oxygen. Once fully sedated, mice were subcutaneously administered dexamethasone (0.025 mg per 100 g; 07-808-8194, Phoenix Pharm, Inc.) to reduce post-surgical inflammation, atropine (0.005 mg per 100 g; 54925-063-10, Med-Pharmex, Inc.) to prevent lung secretions, and ketoprofen (0.5 mg per 100 g; Zoetis, Inc.) to reduce post-surgical inflammation and provide post-surgical analgesia. Mice were then provided with atropine (0.005 mg per 100 g) and 5% glucose in saline (1 ml per 100 g) every hour while anesthetized. The hair was shaved from the back of the neck up to the eyes. Mice were placed on a stereotaxic frame over a feedback-controlled heating blanket to ensure body temperature remained at 37° C. The head was firmly secured and eye ointment was applied to prevent the animal’ s eyes from drying out. The operating area was then sterilized by wiping the skin with iodine and 70% ethanol three times. The mice were given 0.1 ml bupivacaine at the site of incision to serve as a local anesthetic. The skin over the top of the skull was removed and the skull exposed. Using a high-speed drill with different sized bits, a ∼6-mm diameter craniotomy was performed over the cerebral cortex, rostral to the lambda point and caudal to bregma. The exposed brain was then covered with a sterile 8-mm diameter glass coverslip which was glued to the skull surface using cyanoacrylate adhesive. Using dental cement, a small well around the window was created. After completion of the craniotomy, mice were returned to their cages and were subcutaneously administered ketoprofen (0.5 mg per 100 g) and dexamethasone (0.025 mg per 100 g) once daily for three days, and their cages were placed on a heating pad during this time. Animals were given three weeks to recover from the surgery before imaging experiments.

### IN-VIVO TWO-PHOTON MICROSCOPY

For imaging sessions, mice were anesthetized with 3% isoflurane and then maintained at 1.5 – 2% isoflurane in 100% oxygen. Atropine and glucose were provided, as described above. Eye ointment was applied to prevent the eyes from drying out. Mice were placed on a stereotactic frame over a feedback-controlled heating pad to keep body temperature at 37° C. To fluorescently label the microvasculature, Texas Red dextran (50 μl, 2.5% w/v, molecular weight (MW) = 70,000 kDA, Thermo Fisher Scientific) in saline was injected retro-orbitally. Although we did not use these labels for any analysis, these mice also had Rhodamine 6G (0.1 ml, 1 mg/ml in 0.9% saline, Acros Organics, Pure) injected into the bloodstream to label leukocytes and blood platelets and Hoechst 33342 (50 μl, 4.8 mg/ml in 0.9% saline, Thermo Fisher Scientific) to label leukocytes. A custom-built two-photon excitation fluorescence (2PEF) microscope was used to acquire three-dimensional images of the cortical vasculature and to measure red blood cell flow in specific capillaries. Imaging was performed with 830-nm, 75-fs pulses from a Ti-Sapphire laser oscillator (Vision S, Coherent). Lasers were scanned by galvanometric scanners (1 frame/second) and ScanImage software was used to control data acquisition [64]. For obtaining broad maps of the cortical surface vasculature, a 4x magnification air objective (numerical aperture of 0.28, Olympus) was used. For high-resolution imaging, a 25x water-immersion objective lens (numerical aperture of 0.95, Olympus) was used. The emitted fluorescence from Texas Red was detected on a photomultiplier tube through an emission filter with a 641-nm center wavelength and a 75-nm bandwidth. Stacks of images were created by repeatedly taking images axially spaced at 1 μm up to a depth of ∼300 μm. Additionally, centerline line scans and image stacks across the diameter of ∼15 capillaries within the imaging area were obtained in each mouse.

### ANALYSIS OF CAPILLARY BLOOD FLOW AND VESSEL DIAMETER

Blood flow velocities were determined based on the movement of red blood cells (RBCs). Since the injected dye (Texas Red) labels the blood plasma only, RBCs are seen as dark patches within the capillaries. We acquired repetitive line scans along the centerline of individual capillaries, forming space-time images with diagonal streaks due to moving RBCs. As previously described [65], we used a Radon transform-based algorithm to determine the slope of these streaks and thus quantify RBC flow speed. Post-hoc sensitivity analysis (G*Power; α=0.05, β=0.80) with our realized SD and means between running and sedentary groups suggested we could detect a 30% change in flow speed. Capillary diameters were extracted from image stacks taken along with each line scan.

### CROWD-SOURCED SCORING OF CAPILLARIES AS FLOWING OR STALLED

In the 2PEF microscopy used to take three-dimensional image stacks of the vasculature, the intravenously injected fluorescent dye labels the blood plasma, but not red blood cells. Because each capillary segment was visible for multiple frames in the image stack, flowing segments showed different patterns of fluorescent and dark patches in successive frames due to moving blood cells. In capillary segments with stalled blood flow, the dark shadows from non-moving cells remained fixed across all frames where the capillary segment was visible. We used a purpose-built citizen science data analysis platform, StallCatchers.com, that enabled volunteers to score ∼26,000 individual capillary segments as either flowing or not using 2PEF image stacks. We recently reported on the methodological details and validation of this StallCatchers based scoring [58]. Briefly, we used a convolutional neural network, called DeepVess, to segment the 2PEF image stack into voxels that were within vs. outside the vasculature [66]. Individual capillary segments were identified using standard dilation and thinning operations to define vessel centerlines [67], with capillary segments defined as the path between two junctions. To restrict this analysis to capillaries, we excluded all segments with diameter greater than 10 µm. Image stacks were then created that each had a single identified capillary segment outlined and these stacks were analyzed by citizen scientists using StallCatchers to score the identified segment as flowing or stalled. Each segment was scored by multiple volunteers, each of whom had a sensitivity defined by their performance on capillary segments we knew to be flowing or stalled, and we computed a weighted “crowd confidence” score representing the likelihood of each segment being stalled. Laboratory researchers then looked at these capillary segments as a final validation, starting with segments with the highest crowd confidence score for being stalled. The crowd confidence score ranged from 0 to 1, and laboratory researchers examined vessels with a score between 0.5 and 1, a total of 259 capillary segments. For crowd confidence scores between 0.9 and 1, laboratory researchers concluded 95% were stalled, while for crowd confidence scores between 0.5 and 0.6, only 1 out of 80 vessels were not flowing. The initial scoring by citizen scientists decreased the number of capillary segments laboratory researchers needed to evaluate by a factor of 100. We report the density of non-flowing capillaries as stalls per cubic millimeter.

### CHARACTERIZATION OF GEOMETRIC PROPERTIES OF CORTICAL CAPILLARIES

For each identified capillary segment from the 2PEF image stacks that were segmented using DeepVess [66], we calculated the diameter (averaged along the length of the segment), the segment length (distance along the centerline of the vessel between two junctions), and the tortuosity (segment length divided by the Euclidean distance between the two junctions).

### IMMUNOSTAINING OF BRAIN TISSUE

For evaluating cell proliferation, mice received intraperitoneal injection of 5-ethynyl-2’ - deoxyuridine (EdU; E10415, ThermoFisher Scientific) at a dose of 25 mg/kg body weight every day for four days before harvesting the brains. Mice were sacrificed by lethal injection of pentobarbital (5 mg/100 mg). Brains were extracted and cut in half along the center line. One hemisphere was snap frozen in liquid nitrogen for future protein extraction and the other half was kept in 4% paraformaldehyde (PFA) in phosphate buffered saline (PBS) for 48 hours at 4° C and subsequently placed in 30% sucrose for histological analysis. Immunohistochemistry was performed on a set of coronal sections cut on a cryotome with OCT media at a thickness of 30 μm. From each mouse, every sixth section was mounted, washed with PBS, and blocked 1 hour at room temperature (3% goat serum, 0.1% Triton-X100 in PBS). Sections were then incubated overnight at 4°C with primary antibodies against Iba1 (1:500, rabbit anti-mouse Iba1; 019-19741, WAKO) and GFAP (1:500, chicken anti-mouse GFAP; ab53554, Abcam) in the same buffer. Secondary antibodies (goat anti-chicken Alexa Fluor 488, goat anti-rabbit Alexa Fluor 594; ThermoFisher Scientific) were used at 1:300 dilution and added to the slides for 3 hours at room temperature. Cell proliferation was detected through use of the Click-iTTM EdU Cell Proliferation Kit for Imaging (Alexa Fluor 647 dye, ThermoFisher Scientific). Methoxy-X04 was used to counterstain amyloid deposits (1 mg/ml MeO-X04 (5 mg/ml in 10% DMSO, 45% propylene glycol, and 45% saline) for 15 minutes at room temperature, 4920, Tocris). Hoechst 33342 was used to label cell nuclei (3 μg/ml, ThermoFisher Scientific). Sections were washed with PBS before mounting with Richard-Allan Scientific Mounting Medium (4112APG, ThermoFisher Scientific). Images were obtained using confocal microscopy (Zeiss Examiner.D1 AXIO) operated with Zen 1.1.2 software. Z-stack images of the hippocampal and cortical regions of each slide were acquired with 1 μm optical section thickness (three adjacent images between the suprapyramidal blade of the granule cell layer of the dentate gyrus and CA1, one image in CA3, and two adjacent images in the cerebral cortex taken directly toward the cortical surface from the hippocampus). Images were then binarized using a manually determined threshold. Appropriate thresholds varied between mice and were adjusted to ensure that all morphologically relevant objects were recognized. The fraction of pixels above threshold (%Area) and the integrated density (product of mean gray value and area) was determined across sections for cortical and hippocampal regions. All sections were stained and imaged in parallel. Researchers were blinded in this analysis as to whether mice were part of the running or sedentary group.

### ELISA ASSAY

The frozen half-brains were weighed and homogenized in 1 ml PBS containing complete protease inhibitor (Roche Applied Science) and 1 mM AEBSF (Sigma) using a Dounce homogenizer. The homogenates were sonicated for 5 min and centrifuged at 14,000 *g* for 30 min at 4° C. The supernatant (PBS-soluble fraction) was removed and stored at −80° C. The pellet was re-dissolved in 0.5 ml 70% formic acid, sonicated for 5 min, and centrifuged at 14,000 *g* for 30 min at 4° C, and the supernatant was removed and neutralized using 1 M Tris buffer at pH 9 (insoluble fraction). Protein concentration was measured using the Pierce BCA Protein Assay (ThermoFischer Scientific). The extracts were then diluted to equalize different protein concentrations. These samples were analyzed by sandwich ELISA for Aβ1-40 and Aβ1-42 using commercial ELISA kits and following the manufacturer’ s protocol (ThermoFischer Scientific). The Aβ concentration was calculated by comparing the sample absorbance with the absorbance of known concentrations of synthetic Aβ1-40 or Aβ1-42 standards assayed on the same plate. Data was acquired with a Synergy HT plate reader (Biotek) and analyzed using Gen5 software (BioTek) and Prism8 (GraphPad).

### STATISTICAL ANALYSIS

To determine statistical significance of differences between groups, the data was first tested for normality using the Shapiro-Wilk normality test. In case of normality, the statistical comparison was performed using an unpaired t-test for comparison between two groups. In case of non-normal distribution, the Mann-Whitney test (two groups) was used. For the analysis of blood flow speed and capillary diameter, a nested two-way analysis of variance (ANOVA) was carried out to account for possible interactions between individual mice. P-values less than 0.05 were considered statistically significant and we used a standardized set of significance indicators in figures: * *P* < 0.05, ** *P* < 0.01, *** *P* < 0.001, **** *P* < 0.0001. Boxplots show the median with a black line; the mean is indicated with a red line. The box spans between the 25th and the 75th percentile of the data, defined as the interquartile range (IQR). Whiskers of the boxplots extend from the lowest datum within 1.5 times the IQR of the lower quartile of the data to the highest datum within 1.5 times the IQR of the highest quartile of the data. Correlation analysis was performed via the Pearson Product-Moment Correlation, with best fit lines indicated in red. Statistical analysis was performed and graphs were created using Prism8 (GraphPad).

## RESULTS

### RUNNING DISTANCE

To determine the effects of three months of voluntary exercise in 13 month old APP/PS1 mice, we provided single-housed mice with unrestricted access to a monitored running wheel inside their home cages (Fig 1A) [68]. All mice reliably ran on the wheels on a daily basis, with just under a factor of three variance in the average daily running distance between individual animals (Fig 1B). Sedentary, control APP/PS1 mice were singly housed for the same duration, but without access to a running wheel.

**Fig 1.**
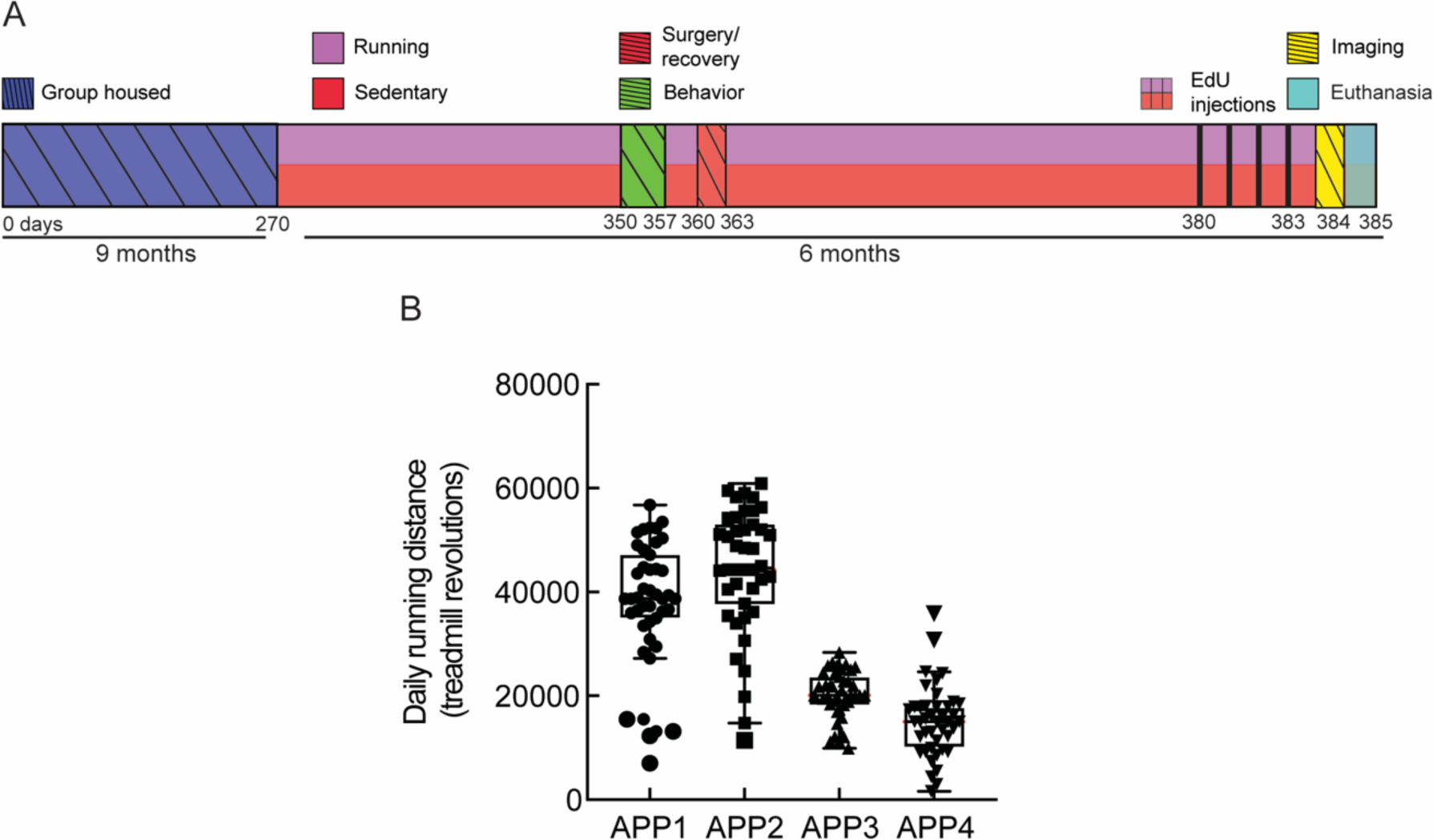
APP/PS1 mice reliably run when given unrestricted access to a running wheel in their home cage. **A**. Timeline of the study. After three months of voluntary exercise on a wheel running (or standard housing for the sedentary control group), mice underwent behavioral testing. Next, a craniotomy was performed, after which mice were allowed three weeks to fully recover. The cortical vasculature was then imaged with in-vivo two photon excitation fluorescence microscopy. Finally, after four consecutive days of EdU-injections, mice were euthanized. Harvested brain tissue was then used for immunohistochemistry. **B**. Each dot is the number of wheel revolutions per day over 45 days for the four APP/PS1 mice in the running group. Data show mean + SD.

### RUNNING IMPROVED SOME MEASURES OF SPATIAL MEMORY PERFORMANCE

To investigate cognitive function in running versus sedentary APP/PS1 mice, we performed a variety of behavioral tests aimed at testing different aspects of memory performance. Running mice achieved a significantly higher preference score in the object replacement test (OR) than sedentary mice, i.e. they spent significantly more time exploring the replaced object than the object in the familiar location as compared to the sedentary mice (Fig 2A; *P* = 0.014). There was also a significant correlation between individual running distances and OR performance (Fig 2B; R^2^ = 0.58 *P* = 0.028). Conversely, we did not observe an increase in spontaneous alternation between the three arms of the Y-maze (Fig 2C), nor increased exploration time for a novel object in the novel object recognition (NOR) test (Fig 2E), nor increased time spent in the center of the arena in an open field (OF) test (Fig. 2G) in running as compared to sedentary APP/PS1 mice. Similarly, none of these measures was significantly correlated with individual running distances (Fig 2D, 2F, and 2H). Additionally, there was no significant difference in time spent exploring objects in the OR test (Fig S1A) or NOR test (Fig S1B), nor in the number of arm entries in the Y-maze (Fig S1C), nor in the average track length during the OF test (Fig S1D) between running and sedentary groups.

**Fig 2.**
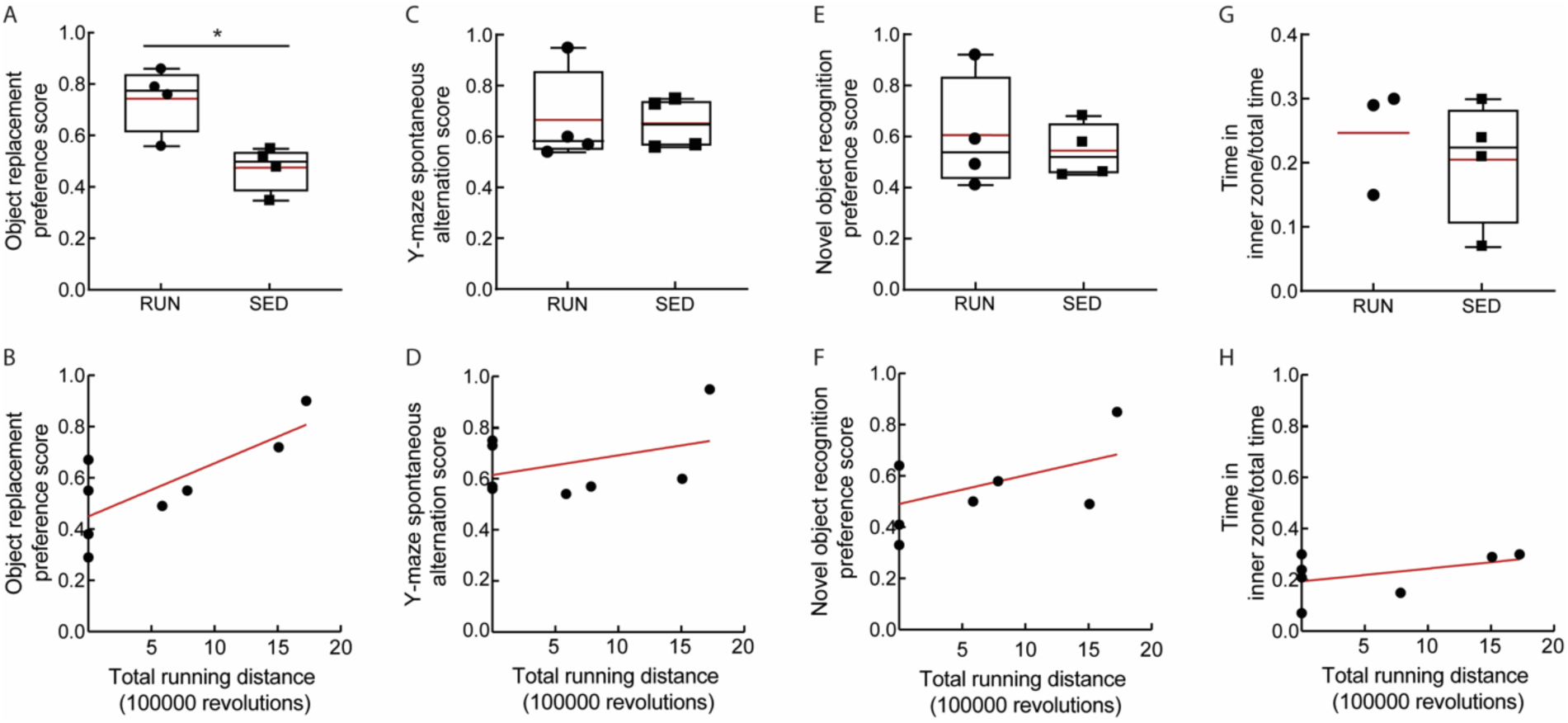
Running improved performance in the object replacement test, but did not show strong effects in other short-term memory tests. All behavioral testing results are shown for running (RUN) and sedentary (SED) APP/PS1 mice. **A** Preference score for OR (P = 0.01). **B** Correlation between OR preference score and total running distance (R2 = 0.6; P = 0.03). **C** Spontaneous alternation score in Y-maze. **D** Correlation between spontaneous alternation and total running distance. **E** Preference score in NOR test. **F** Correlation between NOR preference score and total running distance. **G** Time spent in the arena center in the open field test. One value was excluded in the running group due to a software failure during the test. **H** Correlation between time spent in the arena center and total running distance. Animal numbers: RUN: *n* = 4 (OF: 3); SED: *n* = 4.

### RUNNING INCREASED HIPPOCAMPAL NEUROGENESIS BUT DID NOT DECREASE BRAIN INFLAMMATION NOR AMYLOID DEPOSITION

Exercise has been shown to exert its effects through multiple pathways, with increased hippocampal neurogenesis being one of the most consistently described effects in both patients [69], and in AD mouse models [70, 71]. Consistent with these previous findings, we found an increased number of proliferating neuronal stem cells in the dentate gyrus of running, as compared to sedentary, mice, as detected by the number of EdU-positive cells (Fig 3A and 3B; *P* = 0.03). In the same hippocampal tissue sections, as well as in cortical slices (Fig 3C), we further examined the density of Iba1 and GFAP staining to quantify any differences in the density of microglia and astrocytes, respectively, providing a measure of brain inflammation. We also quantified the density of amyloid plaques that were labeled with Methoxy-X04. We found no notable differences in microglia (Fig 3D) or astrocyte (Fig 3E) density, nor in the number of amyloid deposits (Fig 3F) between running and sedentary APP/PS1 mice in both hippocampal and cortical regions. Using ELISA assays, we quantified the concentration of Aβ1-40 and Aβ1-42 in brain lysates and saw no differences between running and sedentary mice in both the soluble and insoluble fractions (Fig. 4).

**Fig 3.**
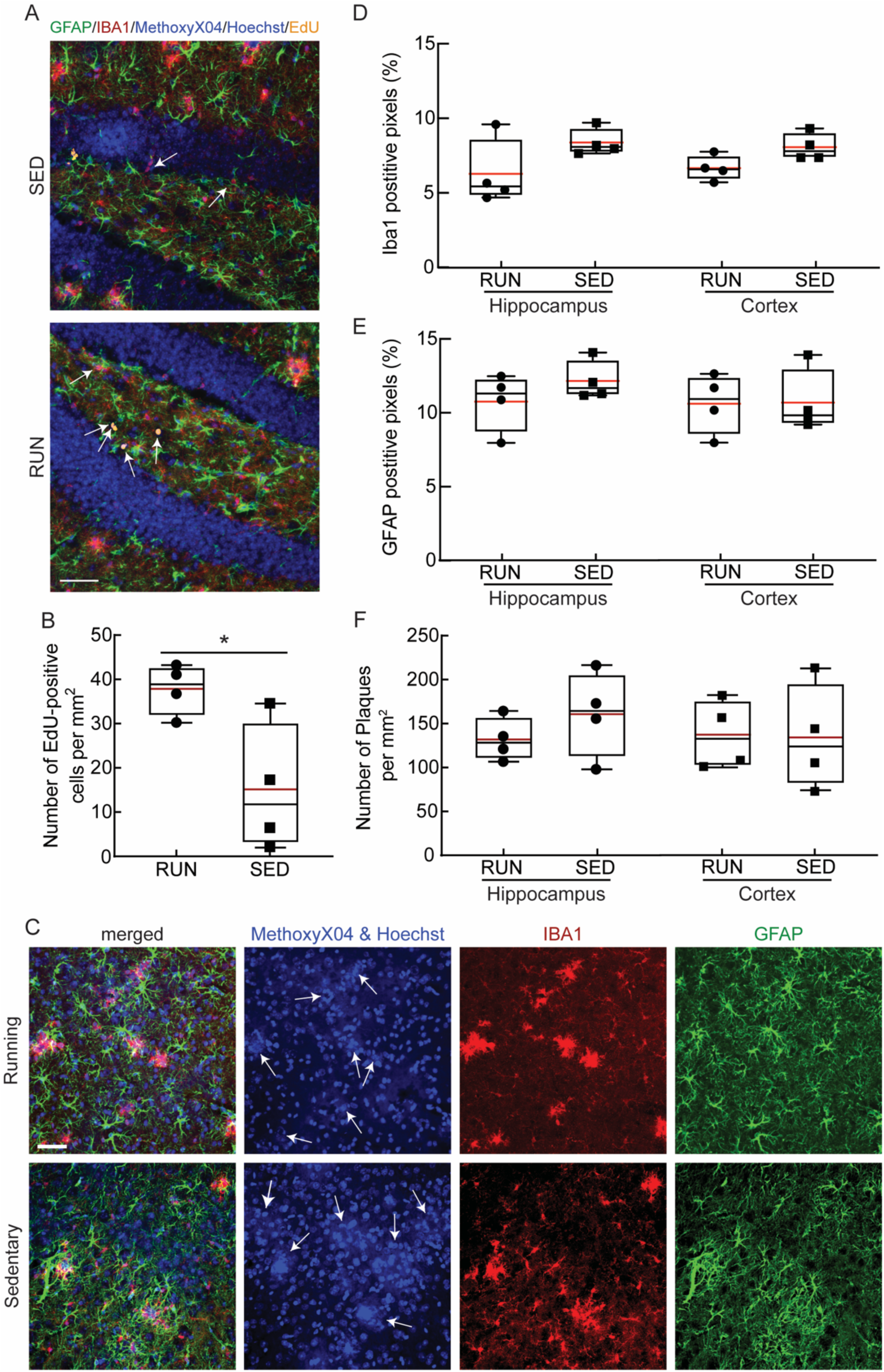
Running increased neural stem cell proliferation in the dentate gyrus, but did not decrease inflammation nor amyloid plaque burden in APP/PS1. **A** Representative confocal images of tissue sections from the hippocampus of sedentary (top) and running (bottom) APP/PS1 mice, with labeling of astrocytes (anti-GFAP, green), microglia (anti-Iba1, red), amyloid plaques (Methoxy-X04, blue; no amyloid plaques visible in these fields), cell nuclei (Hoechst, blue), and proliferating cells (EdU, yellow, indicated with arrows). **B** Boxplot of the density of EdU positive cells in the dentate gyrus from running (RUN) and sedentary (SED) APP/PS1 mice (P = 0.03, Mann-Whitney test). **C** Representative confocal images of tissue sections from the cortex from running (top) and sedentary (bottom) mice. Labeling is the same as in panel A. Images to the right show individual channels. The white arrows in the second column of images indicate Methoxy-X04 labeled amyloid plaques, which are distinguished from Hoechst-labeled cell nuclei by morphological differences. **D** Microglial density, represented as the fractional area that is Iba-1 positive; **E** Astrocyte density, represented as the fractional area that was GFAP positive; and **F** the number of Methoxy-X04 positive plaques per unit area, from the hippocampus and cortex of RUN and SED APP/PS1 mice. Animal numbers: RUN: *n* = 4; SED: *n* = 4. Scale bars indicates 50 μm.

**Fig 4.**
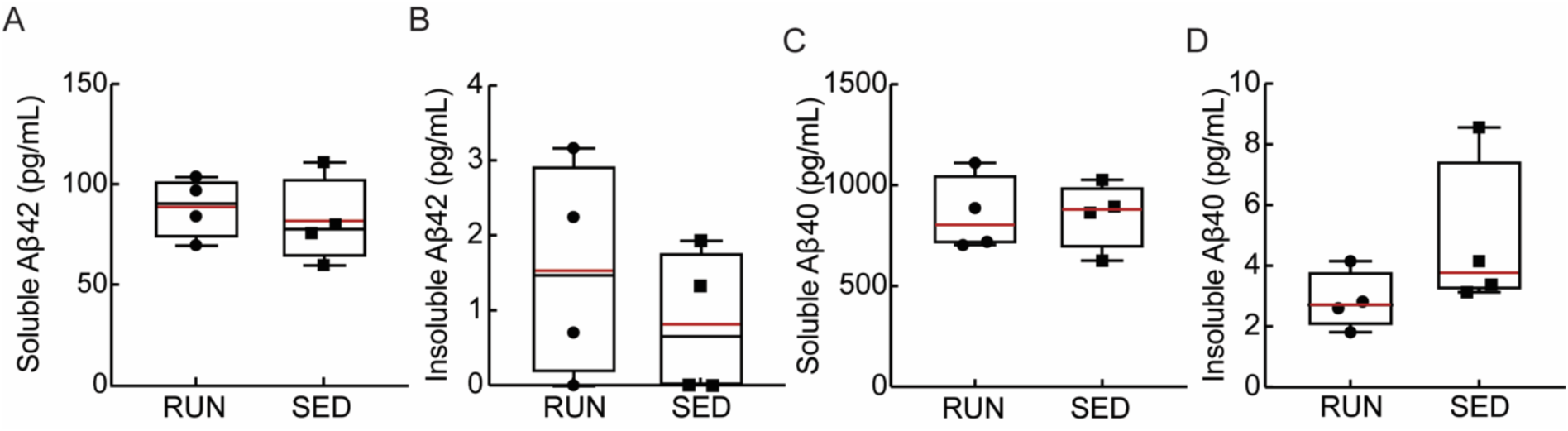
Running did not alter the levels of soluble and insoluble amyloid beta monomers in the brain. Soluble (A and C) and insoluble (B and D) concentrations of Aβ1-42 (A and B) and Aβ1-40 (C and D) in brain lysates from running (RUN) and sedentary (SED) APP/PS1 mice, as measured using ELISA assays.

### VOLUNTARY RUNNING DID NOT AFFECT CAPILLARY BLOOD FLOW

The trends toward improved cognitive function (Fig 2), the clear increase in neurogenesis, as well as the lack of a notable impact on brain inflammation and amyloid deposition (Fig. 3 and 4) that we observed are consistent with previous studies of the impact of exercise on mouse models of AD [72]. We next sought to determine if these exercise-mediated changes were correlated with an increase in brain blood flow, perhaps linked to a decrease in the incidence of non-flowing capillaries. We used a crowd-sourced approach to score individual capillary segments as flowing or stalled based on the motion of red blood cells (which appear as dark patches within the fluorescently-labeled blood plasma in 2PEF image stacks) (Fig 5A). Contrary to our initial hypothesis, we did not observe a decrease in the incidence of stalled capillaries in running APP/PS1 mice as compared to sedentary controls (Fig 5B). We further quantified red blood cell flow speed and vessel diameter in cortical capillaries (Fig. 5C), and saw no differences, on average, between running and sedentary APP/PS1 mice (Fig 5D). Capillary speed and diameter were also not found to be correlated with the total distance ran by animals (Fig S2). To further explore potential exercise-induced changes in the brain vasculature, we used a convolutional neural network-based segmentation algorithm, DeepVess [66], to segment 3D image stacks of the cortical vasculature of running (Fig 6A) and sedentary (Fig 6B) APP/PS1 mice and we characterized the capillary density (Fig 6C), capillary diameter (Fig 6D), capillary segment length (Fig 6E), and capillary tortuosity (Fig 6F). None of these parameters differed between running and sedentary APP/PS1 mice.

**Fig 5.**
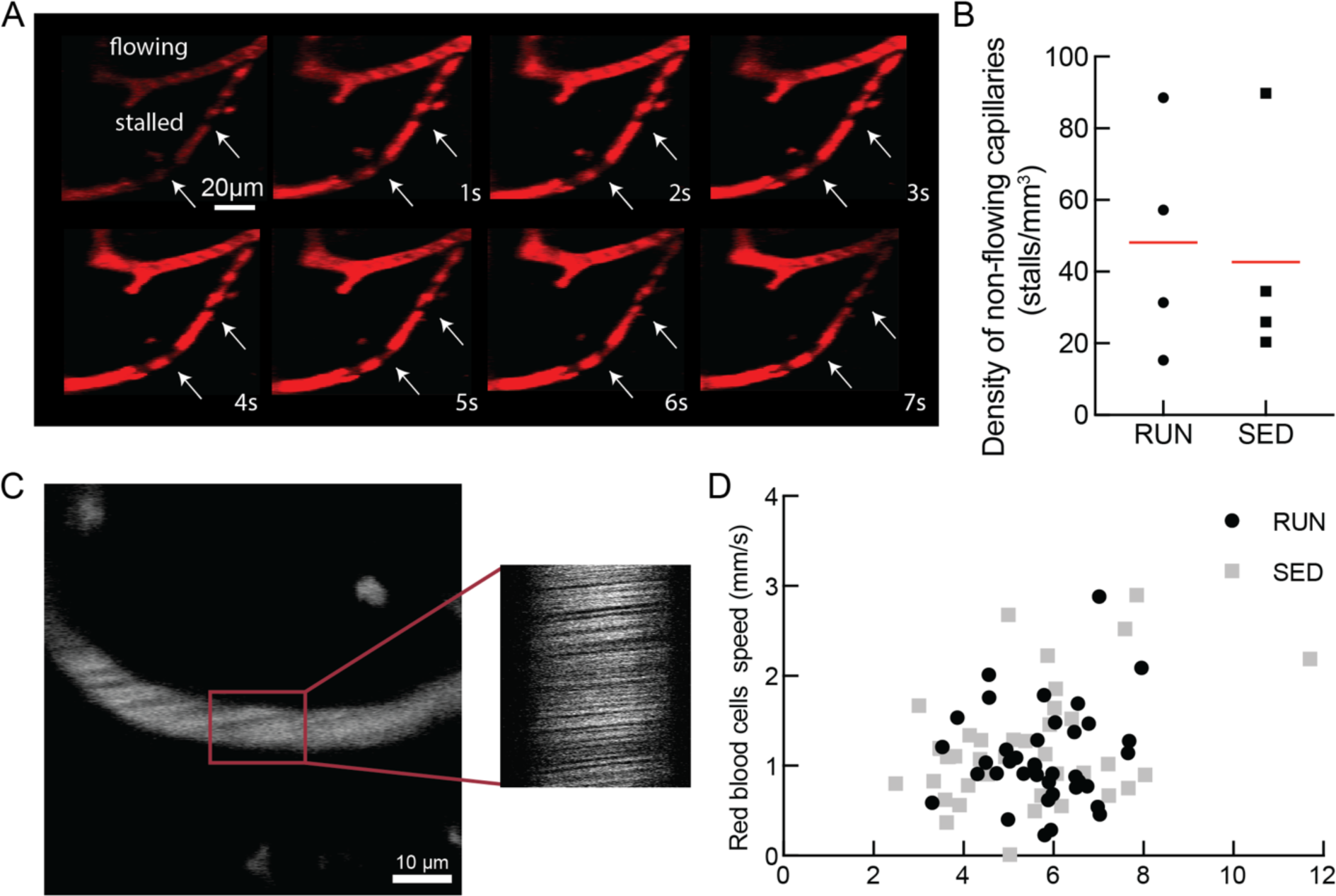
Running did not decrease the incidence of non-flowing capillaries, not increase capillary blood flow in the brain of APP/PS1 mice. **A** Representative 2PEF image sequence showing both flowing and stalled capillary segments over 7 s. The blood plasma was labeled with Texas Red-dextran and the dark patches in the vessel lumen were formed by blood cells. The lack of motion of these dark patches in the lower capillary indicates stalled blood flow. (Stalled capillary is indicated with arrows). **B** Density of capillaries with stalled blood flow in running (RUN) and sedentary (SED) APP/PS1 mice. **C** Representative 2PEF image of a cortical capillary (left) and a space-time image from repetitive line scans along the centerline of the capillary segment (right). The diagonal streaks in the space-time image are formed by moving red blood cells, with a slope that is inversely proportional to the blood flow speed. **D** Capillary blood flow speed plotted as a function of vessel diameter for cortical capillaries from running (RUN) and sedentary (SED) APP/PS1 mice.

**Fig. 6.**
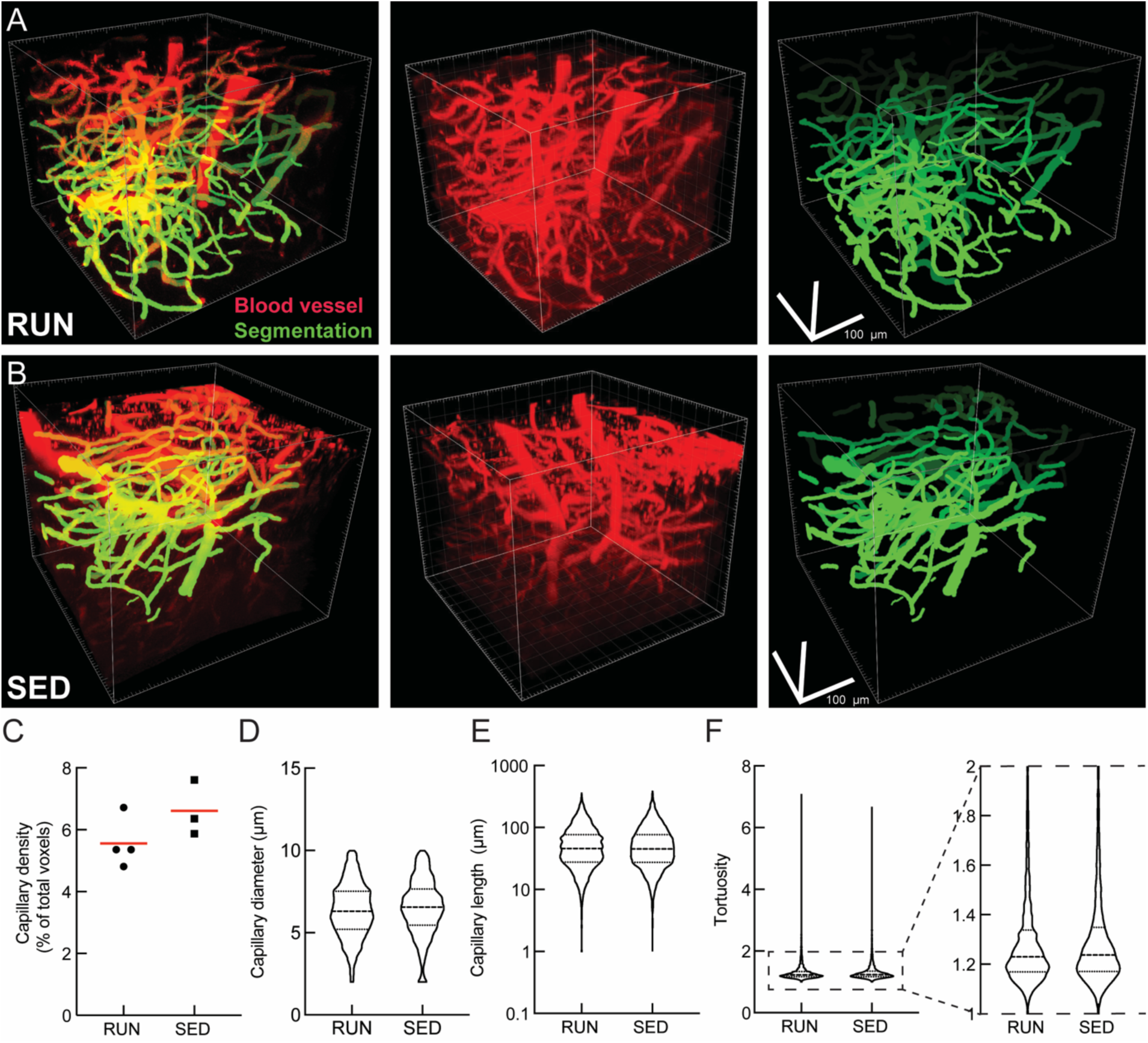
Running did not alter capillary density or geometry. **A** and **B** Representative 2PEF image stacks of fluorescently-labeled cortical vasculature (middle) and the segmentation of this image (right) from running (RUN) and sedentary (SED) APP/PS1 mice, respectively. **C** Density; **D** Diameter; **E** Length and; **F** Tortuosity of capillary segments from running (RUN; n = 5514 capillaries) and sedentary (SED; n = 7251 capillaries) APP/PS1 mice.

## DISCUSSION

Physical activity is correlated with attenuation of cognitive impairment, amelioration of age-related changes in the brain, and reduced risk of dementia [73, 74]. While exercise has been found to have a positive effect on cognition and brain health in aging individuals [75], the underlying mechanisms have not been fully elucidated. In this study, we investigated the effects of several months of voluntary wheel running on memory function, AD-related brain pathology, and cortical blood flow in aged APP/PS1 mice.

Impairments in spatial learning and memory in APP/PS1 mice have been reported in mice as young as seven months of age [76, 77]. We found performance in the object replacement task in 12-month-old APP/PS1 mice was markedly improved by three months of voluntary wheel running, as compared to sedentary controls, while no improvements were detected in the novel object recognition task and Y-maze tasks. This exercise-related improvement in performance on some memory-related tasks but not on others is consistent with similarly mixed impacts of exercise in previous studies of memory function in mouse models of AD [72] and in AD patients [78]. While a variety of brain regions are likely involved in each of these memory tasks, it has been shown that memory of an object’ s spatial location (evaluated with object replacement task) is highly dependent on hippocampal regions, while memory of an object’ s intrinsic characteristics (evaluated with novel object task) also involves significant contributions from other brain regions, such as the temporal lobe [79-81]. It is possible that exercise differentially improves function in different brain regions and this contributes to the mixed impact of exercise on different memory tasks.

Consistent with a broad consensus in the literature [37, 40], we observed increased neural stem cell proliferation in the dentate gyrus of the hippocampus in exercising APP/PS1 mice, as compared to sedentary controls. Exercise did not, however, lead to changes in brain inflammation, as assayed by microglia and astrocyte density, or in amyloid pathology, as assayed by amyloid plaque density and amyloid-beta monomer concentrations. A recent study in the 5xFAD mouse model of AD similarly found no decreases in measures of brain inflammation or in the density of amyloid deposits due to exercise [82].

We have shown that neutrophils plug a small fraction of brain capillary segments in the APP/PS1 and 5xFAD mouse models of AD, and that this contributes to the overall brain blood flow reductions seen in these mice [57]. Contrary to our initial hypothesis that exercise could decrease the microvascular dysfunction that underlies these capillary stalls, we did not find a decrease in the number of non-flowing capillaries in running APP/PS1 mice, as compared to sedentary controls. Further, we did not detect any differences in average capillary blood flow speeds between running and sedentary APP/PS1 mice. While there is clear consensus that reduced brain blood flow in both AD patients and mouse models is a key feature of the disease, the effects of exercise on cortical blood flow in AD remain up for debate. Consistent with our findings in APP/PS1 mice, however, recent studies in patients with dementia (diagnosed as mild to moderate AD in Refs. [83] and [84]) showed that 3 times per week 60 min of aerobic exercise led to improved cognition [83, 85, 86], but this effect was not correlated with increased cerebral blood flow [84, 87]. Finally, we did not observe changes in the density or geometry of cortical capillaries in APP/PS1 mice that exercised, compared to sedentary controls.

While we did not observe an impact of exercise on the capillary stalling phenomena we recently tied to brain blood flow deficits in AD mouse models, there are other aspects of cortical microvascular dysfunction that have been shown to occur in mouse models of AD that we did not examine. For example, mouse models of AD have shown increased blood brain barrier permeability [88], more contractile pericytes around capillaries [54], attenuation of cerebral blood flow regulation mechanisms (neurovascular coupling [89] and autoregulation [90]), as well as hypercoaguability [55, 56]. It is possible that exercise could positively impact some of these other aspects of microvascular dysfunction in AD, although our observation of no differences in average capillary flow speeds between exercising and sedentary APP/PS1 mice suggests that pathologies that impact flow are not likely modulated by exercise.

## ACKNOWLEDGMENTS

The confocal microscopy imaging data in this manuscript was acquired through the Cornell University Biotechnology Resource Center, with National Institutes of Health grant S10RR025502 funding for the shared Zeiss LSM 710 Confocal.

## REFERENCES

1. Mattson MP. Pathways towards and away from Alzheimer’s disease. Nature. 2004;430(7000):631-9. Epub 2004/08/06. doi: 10.1038/nature02621. PubMed PMID: 15295589; PubMed Central PMCID: PMCPMC3091392.

2. Yaffe K, Barnes D, Nevitt M, Lui LY, Covinsky K. A prospective study of physical activity and cognitive decline in elderly women: women who walk. Arch Intern Med. 2001;161(14):1703-8. Epub 2001/08/04. PubMed PMID: 11485502.

3. Barnes DE, Yaffe K, Satariano WA, Tager IB. A longitudinal study of cardiorespiratory fitness and cognitive function in healthy older adults. J Am Geriatr Soc. 2003;51(4):459-65. Epub 2003/03/27. doi: 10.1046/j.1532-5415.2003.51153.x. PubMed PMID: 12657064.

4. Rovio S, Kareholt I, Helkala EL, Viitanen M, Winblad B, Tuomilehto J, et al. Leisure-time physical activity at midlife and the risk of dementia and Alzheimer’s disease. Lancet Neurol. 2005;4(11):705-11. Epub 2005/10/22. doi: 10.1016/S1474-4422(05)70198-8. PubMed PMID: 16239176.

5. Larson EB, Wang L, Bowen JD, McCormick WC, Teri L, Crane P, et al. Exercise is associated with reduced risk for incident dementia among persons 65 years of age and older. Ann Intern Med. 2006;144(2):73-81. Epub 2006/01/19. doi: 10.7326/0003-4819-144-2-200601170-00004. PubMed PMID: 16418406.

6. Kramer AF, Hahn S, Cohen NJ, Banich MT, McAuley E, Harrison CR, et al. Ageing, fitness and neurocognitive function. Nature. 1999;400(6743):418-9. Epub 1999/08/10. doi: 10.1038/22682. PubMed PMID: 10440369.

7. Anderson-Hanley C, Arciero PJ, Brickman AM, Nimon JP, Okuma N, Westen SC, et al. Exergaming and older adult cognition: a cluster randomized clinical trial. Am J Prev Med. 2012;42(2):109-19. Epub 2012/01/21. doi: 10.1016/j.amepre.2011.10.016. PubMed PMID: 22261206.

8. Lautenschlager NT, Cox KL, Flicker L, Foster JK, van Bockxmeer FM, Xiao J, et al. Effect of physical activity on cognitive function in older adults at risk for Alzheimer disease: a randomized trial. JAMA. 2008;300(9):1027-37. Epub 2008/09/05. doi: 10.1001/jama.300.9.1027. PubMed PMID: 18768414.

9. Nagamatsu LS, Chan A, Davis JC, Beattie BL, Graf P, Voss MW, et al. Physical activity improves verbal and spatial memory in older adults with probable mild cognitive impairment: a 6-month randomized controlled trial. J Aging Res. 2013;2013:861893. Epub 2013/03/20. doi: 10.1155/2013/861893. PubMed PMID: 23509628; PubMed Central PMCID: PMCPMC3595715.

10. Garcia-Mesa Y, Lopez-Ramos JC, Gimenez-Llort L, Revilla S, Guerra R, Gruart A, et al. Physical exercise protects against Alzheimer’s disease in 3xTg-AD mice. J Alzheimers Dis. 2011;24(3):421-54. Epub 2011/02/08. doi: 10.3233/JAD-2011-101635. PubMed PMID: 21297257.

11. Bo H, Kang W, Jiang N, Wang X, Zhang Y, Ji LL. Exercise-induced neuroprotection of hippocampus in APP/PS1 transgenic mice via upregulation of mitochondrial 8-oxoguanine DNA glycosylase. Oxid Med Cell Longev. 2014;2014:834502. Epub 2014/12/30. doi: 10.1155/2014/834502. PubMed PMID: 25538817; PubMed Central PMCID: PMCPMC4236906.

12. Revilla S, Sunol C, Garcia-Mesa Y, Gimenez-Llort L, Sanfeliu C, Cristofol R. Physical exercise improves synaptic dysfunction and recovers the loss of survival factors in 3xTg-AD mouse brain. Neuropharmacology. 2014;81:55-63. Epub 2014/02/04. doi: 10.1016/j.neuropharm.2014.01.037. PubMed PMID: 24486380.

13. Tapia-Rojas C, Aranguiz F, Varela-Nallar L, Inestrosa NC. Voluntary Running Attenuates Memory Loss, Decreases Neuropathological Changes and Induces Neurogenesis in a Mouse Model of Alzheimer’s Disease. Brain Pathol. 2016;26(1):62-74. Epub 2015/03/13. doi: 10.1111/bpa.12255. PubMed PMID: 25763997.

14. Lourenco MV, Frozza RL, de Freitas GB, Zhang H, Kincheski GC, Ribeiro FC, et al. Exercise-linked FNDC5/irisin rescues synaptic plasticity and memory defects in Alzheimer’s models. Nat Med. 2019;25(1):165-75. Epub 2019/01/09. doi: 10.1038/s41591-018-0275-4. PubMed PMID: 30617325; PubMed Central PMCID: PMCPMC6327967.

15. Yuede CM, Zimmerman SD, Dong H, Kling MJ, Bero AW, Holtzman DM, et al. Effects of voluntary and forced exercise on plaque deposition, hippocampal volume, and behavior in the Tg2576 mouse model of Alzheimer’s disease. Neurobiol Dis. 2009;35(3):426-32. Epub 2009/06/16. doi: 10.1016/j.nbd.2009.06.002. PubMed PMID: 19524672; PubMed Central PMCID: PMCPMC2745233.

16. Zhao G, Liu HL, Zhang H, Tong XJ. Treadmill exercise enhances synaptic plasticity, but does not alter beta-amyloid deposition in hippocampi of aged APP/PS1 transgenic mice. Neuroscience. 2015;298:357-66. Epub 2015/04/29. doi: 10.1016/j.neuroscience.2015.04.038. PubMed PMID: 25917310.

17. Cho J, Shin MK, Kim D, Lee I, Kim S, Kang H. Treadmill Running Reverses Cognitive Declines due to Alzheimer Disease. Med Sci Sports Exerc. 2015;47(9):1814-24. Epub 2015/01/13. doi: 10.1249/MSS.0000000000000612. PubMed PMID: 25574797.

18. Adlard PA, Perreau VM, Pop V, Cotman CW. Voluntary exercise decreases amyloid load in a transgenic model of Alzheimer’s disease. J Neurosci. 2005;25(17):4217-21. Epub 2005/04/29. doi: 10.1523/JNEUROSCI.0496-05.2005. PubMed PMID: 15858047.

19. Herring A, Munster Y, Metzdorf J, Bolczek B, Krussel S, Krieter D, et al. Late running is not too late against Alzheimer’s pathology. Neurobiol Dis. 2016;94:44-54. Epub 2016/06/18. doi: 10.1016/j.nbd.2016.06.003. PubMed PMID: 27312772.

20. Moore KM, Girens RE, Larson SK, Jones MR, Restivo JL, Holtzman DM, et al. A spectrum of exercise training reduces soluble Abeta in a dose-dependent manner in a mouse model of Alzheimer’s disease. Neurobiol Dis. 2016;85:218-24. Epub 2015/11/14. doi: 10.1016/j.nbd.2015.11.004. PubMed PMID: 26563933.

21. He XF, Liu DX, Zhang Q, Liang FY, Dai GY, Zeng JS, et al. Voluntary Exercise Promotes Glymphatic Clearance of Amyloid Beta and Reduces the Activation of Astrocytes and Microglia in Aged Mice. Front Mol Neurosci. 2017;10:144. Epub 2017/06/06. doi: 10.3389/fnmol.2017.00144. PubMed PMID: 28579942; PubMed Central PMCID: PMCPMC5437122.

22. Nichol KE, Poon WW, Parachikova AI, Cribbs DH, Glabe CG, Cotman CW. Exercise alters the immune profile in Tg2576 Alzheimer mice toward a response coincident with improved cognitive performance and decreased amyloid. J Neuroinflammation. 2008;5:13. Epub 2008/04/11. doi: 10.1186/1742-2094-5-13. PubMed PMID: 18400101; PubMed Central PMCID: PMCPMC2329612.

23. Zhang XL, Zhao N, Xu B, Chen XH, Li TJ. Treadmill exercise inhibits amyloid-beta generation in the hippocampus of APP/PS1 transgenic mice by reducing cholesterol-mediated lipid raft formation. Neuroreport. 2019;30(7):498-503. Epub 2019/03/19. doi: 10.1097/WNR.0000000000001230. PubMed PMID: 30882716.

24. Koo JH, Kang EB, Oh YS, Yang DS, Cho JY. Treadmill exercise decreases amyloid-beta burden possibly via activation of SIRT-1 signaling in a mouse model of Alzheimer’s disease. Exp Neurol. 2017;288:142-52. Epub 2016/11/28. doi: 10.1016/j.expneurol.2016.11.014. PubMed PMID: 27889467.

25. Zhao N, Zhang X, Song C, Yang Y, He B, Xu B. The effects of treadmill exercise on autophagy in hippocampus of APP/PS1 transgenic mice. Neuroreport. 2018;29(10):819-25. Epub 2018/04/20. doi: 10.1097/WNR.0000000000001038. PubMed PMID: 29672446; PubMed Central PMCID: PMCPMC5999367.

26. Yan QW, Zhao N, Xia J, Li BX, Yin LY. Effects of treadmill exercise on mitochondrial fusion and fission in the hippocampus of APP/PS1 mice. Neurosci Lett. 2019;701:84-91. Epub 2019/02/24. doi: 10.1016/j.neulet.2019.02.030. PubMed PMID: 30796962.

27. Zhang L, Chao FL, Luo YM, Xiao Q, Jiang L, Zhou CN, et al. Exercise Prevents Cognitive Function Decline and Demyelination in the White Matter of APP/PS1 Transgenic AD Mice. Curr Alzheimer Res. 2017;14(6):645-55. Epub 2016/12/17. doi: 10.2174/1567205014666161213121353. PubMed PMID: 27978791.

28. Zhou CN, Chao FL, Zhang Y, Jiang L, Zhang L, Luo YM, et al. Sex Differences in the White Matter and Myelinated Fibers of APP/PS1 Mice and the Effects of Running Exercise on the Sex Differences of AD Mice. Front Aging Neurosci. 2018;10:243. Epub 2018/09/04. doi: 10.3389/fnagi.2018.00243. PubMed PMID: 30174598; PubMed Central PMCID: PMCPMC6107833.

29. Chao F, Zhang L, Luo Y, Xiao Q, Lv F, He Q, et al. Running Exercise Reduces Myelinated Fiber Loss in the Dentate Gyrus of the Hippocampus in APP/PS1 Transgenic Mice. Curr Alzheimer Res. 2015;12(4):377-83. Epub 2015/03/31. PubMed PMID: 25817255.

30. Chao FL, Zhang L, Zhang Y, Zhou CN, Jiang L, Xiao Q, et al. Running exercise protects against myelin breakdown in the absence of neurogenesis in the hippocampus of AD mice. Brain Res. 2018;1684:50-9. Epub 2018/01/11. doi: 10.1016/j.brainres.2018.01.007. PubMed PMID: 29317290.

31. Kang EB, Kwon IS, Koo JH, Kim EJ, Kim CH, Lee J, et al. Treadmill exercise represses neuronal cell death and inflammation during Abeta-induced ER stress by regulating unfolded protein response in aged presenilin 2 mutant mice. Apoptosis. 2013;18(11):1332-47. Epub 2013/08/03. doi: 10.1007/s10495-013-0884-9. PubMed PMID: 23907580.

32. Do K, Laing BT, Landry T, Bunner W, Mersaud N, Matsubara T, et al. The effects of exercise on hypothalamic neurodegeneration of Alzheimer’s disease mouse model. PLoS One. 2018;13(1):e0190205. Epub 2018/01/03. doi: 10.1371/journal.pone.0190205. PubMed PMID: 29293568; PubMed Central PMCID: PMCPMC5749759.

33. Sun LN, Qi JS, Gao R. Physical exercise reserved amyloid-beta induced brain dysfunctions by regulating hippocampal neurogenesis and inflammatory response via MAPK signaling. Brain Res. 2018;1697:1-9. Epub 2018/05/08. doi: 10.1016/j.brainres.2018.04.040. PubMed PMID: 29729254.

34. Zhang J, Guo Y, Wang Y, Song L, Zhang R, Du Y. Long-term treadmill exercise attenuates Abeta burdens and astrocyte activation in APP/PS1 mouse model of Alzheimer’s disease. Neurosci Lett. 2018;666:70-7. Epub 2017/12/17. doi: 10.1016/j.neulet.2017.12.025. PubMed PMID: 29246793.

35. Garcia-Mesa Y, Colie S, Corpas R, Cristofol R, Comellas F, Nebreda AR, et al. Oxidative Stress Is a Central Target for Physical Exercise Neuroprotection Against Pathological Brain Aging. J Gerontol A Biol Sci Med Sci. 2016;71(1):40-9. Epub 2015/02/28. doi: 10.1093/gerona/glv005. PubMed PMID: 25720862.

36. Um HS, Kang EB, Koo JH, Kim HT, Jin L, Kim EJ, et al. Treadmill exercise represses neuronal cell death in an aged transgenic mouse model of Alzheimer’s disease. Neurosci Res. 2011;69(2):161-73. Epub 2010/10/26. doi: 10.1016/j.neures.2010.10.004. PubMed PMID: 20969897.

37. van Praag H, Kempermann G, Gage FH. Running increases cell proliferation and neurogenesis in the adult mouse dentate gyrus. Nat Neurosci. 1999;2(3):266-70. Epub 1999/04/09. doi: 10.1038/6368. PubMed PMID: 10195220.

38. Koo JH, Kwon IS, Kang EB, Lee CK, Lee NH, Kwon MG, et al. Neuroprotective effects of treadmill exercise on BDNF and PI3-K/Akt signaling pathway in the cortex of transgenic mice model of Alzheimer’s disease. J Exerc Nutrition Biochem. 2013;17(4):151-60. Epub 2015/01/08. doi: 10.5717/jenb.2013.17.4.151. PubMed PMID: 25566426; PubMed Central PMCID: PMCPMC4241914.

39. Xiong JY, Li SC, Sun YX, Zhang XS, Dong ZZ, Zhong P, et al. Long-term treadmill exercise improves spatial memory of male APPswe/PS1dE9 mice by regulation of BDNF expression and microglia activation. Biol Sport. 2015;32(4):295-300. Epub 2015/12/19. doi: 10.5604/20831862.1163692. PubMed PMID: 26681831; PubMed Central PMCID: PMCPMC4672160.

40. Choi SH, Bylykbashi E, Chatila ZK, Lee SW, Pulli B, Clemenson GD, et al. Combined adult neurogenesis and BDNF mimic exercise effects on cognition in an Alzheimer’s mouse model. Science. 2018;361(6406). Epub 2018/09/08. doi: 10.1126/science.aan8821. PubMed PMID: 30190379; PubMed Central PMCID: PMCPMC6149542.

41. Santos CY, Snyder PJ, Wu WC, Zhang M, Echeverria A, Alber J. Pathophysiologic relationship between Alzheimer’s disease, cerebrovascular disease, and cardiovascular risk: A review and synthesis. Alzheimers Dement (Amst). 2017;7:69-87. Epub 2017/03/10. doi: 10.1016/j.dadm.2017.01.005. PubMed PMID: 28275702; PubMed Central PMCID: PMCPMC5328683.

42. Wiesmann M, Zerbi V, Jansen D, Lutjohann D, Veltien A, Heerschap A, et al. Hypertension, cerebrovascular impairment, and cognitive decline in aged AbetaPP/PS1 mice. Theranostics. 2017;7(5):1277-89. Epub 2017/04/25. doi: 10.7150/thno.18509. PubMed PMID: 28435465; PubMed Central PMCID: PMCPMC5399593.

43. Dai W, Lopez OL, Carmichael OT, Becker JT, Kuller LH, Gach HM. Mild cognitive impairment and alzheimer disease: patterns of altered cerebral blood flow at MR imaging. Radiology. 2009;250(3):856-66. Epub 2009/01/24. doi: 10.1148/radiol.2503080751. PubMed PMID: 19164119; PubMed Central PMCID: PMCPMC2680168.

44. Bennett RE, Robbins AB, Hu M, Cao X, Betensky RA, Clark T, et al. Tau induces blood vessel abnormalities and angiogenesis-related gene expression in P301L transgenic mice and human Alzheimer’s disease. Proc Natl Acad Sci U S A. 2018;115(6):E1289-E98. Epub 2018/01/24. doi: 10.1073/pnas.1710329115. PubMed PMID: 29358399; PubMed Central PMCID: PMCPMC5819390.

45. Decker Y, Muller A, Nemeth E, Schulz-Schaeffer WJ, Fatar M, Menger MD, et al. Analysis of the vasculature by immunohistochemistry in paraffin-embedded brains. Brain Struct Funct. 2018;223(2):1001-15. Epub 2017/12/21. doi: 10.1007/s00429-017-1595-8. PubMed PMID: 29260371.

46. Gutierrez-Jimenez E, Angleys H, Rasmussen PM, West MJ, Catalini L, Iversen NK, et al. Disturbances in the control of capillary flow in an aged APP(swe)/PS1DeltaE9 model of Alzheimer’s disease. Neurobiol Aging. 2018;62:82-94. Epub 2017/11/14. doi: 10.1016/j.neurobiolaging.2017.10.006. PubMed PMID: 29131981.

47. Kisler K, Nelson AR, Rege SV, Ramanathan A, Wang Y, Ahuja A, et al. Pericyte degeneration leads to neurovascular uncoupling and limits oxygen supply to brain. Nat Neurosci. 2017;20(3):406-16. Epub 2017/01/31. doi: 10.1038/nn.4489. PubMed PMID: 28135240; PubMed Central PMCID: PMCPMC5323291.

48. Bannai T, Mano T, Chen X, Ohtomo G, Ohtomo R, Tsuchida T, et al. Chronic cerebral hypoperfusion shifts the equilibrium of amyloid beta oligomers to aggregation-prone species with higher molecular weight. Sci Rep. 2019;9(1):2827. Epub 2019/02/28. doi: 10.1038/s41598-019-39494-7. PubMed PMID: 30808940; PubMed Central PMCID: PMCPMC6391466.

49. Ni R, Rudin M, Klohs J. Cortical hypoperfusion and reduced cerebral metabolic rate of oxygen in the arcAbeta mouse model of Alzheimer’s disease. Photoacoustics. 2018;10:38-47. Epub 2018/04/24. doi: 10.1016/j.pacs.2018.04.001. PubMed PMID: 29682448; PubMed Central PMCID: PMCPMC5909030.

50. Montagne A, Nikolakopoulou AM, Zhao Z, Sagare AP, Si G, Lazic D, et al. Pericyte degeneration causes white matter dysfunction in the mouse central nervous system. Nat Med. 2018;24(3):326-37. Epub 2018/02/06. doi: 10.1038/nm.4482. PubMed PMID: 29400711; PubMed Central PMCID: PMCPMC5840035.

51. Tachibana M, Yamazaki Y, Liu CC, Bu G, Kanekiyo T. Pericyte implantation in the brain enhances cerebral blood flow and reduces amyloid-beta pathology in amyloid model mice. Exp Neurol. 2018;300:13-21. Epub 2017/11/07. doi: 10.1016/j.expneurol.2017.10.023. PubMed PMID: 29106980; PubMed Central PMCID: PMCPMC5745278.

52. Hamel E. Cerebral circulation: function and dysfunction in Alzheimer’s disease. J Cardiovasc Pharmacol. 2015;65(4):317-24. Epub 2014/11/11. doi: 10.1097/FJC.0000000000000177. PubMed PMID: 25384195.

53. Eguchi K, Shindo T, Ito K, Ogata T, Kurosawa R, Kagaya Y, et al. Whole-brain low-intensity pulsed ultrasound therapy markedly improves cognitive dysfunctions in mouse models of dementia - Crucial roles of endothelial nitric oxide synthase. Brain Stimul. 2018;11(5):959-73. Epub 2018/06/03. doi: 10.1016/j.brs.2018.05.012. PubMed PMID: 29857968.

54. Nortley R, Korte N, Izquierdo P, Hirunpattarasilp C, Mishra A, Jaunmuktane Z, et al. Amyloid beta oligomers constrict human capillaries in Alzheimer’s disease via signaling to pericytes. Science. 2019;365(6450). Epub 2019/06/22. doi: 10.1126/science.aav9518. PubMed PMID: 31221773; PubMed Central PMCID: PMCPMC6658218.

55. Cortes-Canteli M, Kruyer A, Fernandez-Nueda I, Marcos-Diaz A, Ceron C, Richards AT, et al. Long-Term Dabigatran Treatment Delays Alzheimer’s Disease Pathogenesis in the TgCRND8 Mouse Model. J Am Coll Cardiol. 2019;74(15):1910-23. Epub 2019/10/12. doi: 10.1016/j.jacc.2019.07.081. PubMed PMID: 31601371; PubMed Central PMCID: PMCPMC6822166.

56. Ahn HJ, Zamolodchikov D, Cortes-Canteli M, Norris EH, Glickman JF, Strickland S. Alzheimer’s disease peptide beta-amyloid interacts with fibrinogen and induces its oligomerization. Proc Natl Acad Sci U S A. 2010;107(50):21812-7. Epub 2010/11/26. doi: 10.1073/pnas.1010373107. PubMed PMID: 21098282; PubMed Central PMCID: PMCPMC3003082.

57. Cruz Hernandez JC, Bracko O, Kersbergen CJ, Muse V, Haft-Javaherian M, Berg M, et al. Neutrophil adhesion in brain capillaries reduces cortical blood flow and impairs memory function in Alzheimer’s disease mouse models. Nat Neurosci. 2019;22(3):413-20. Epub 2019/02/12. doi: 10.1038/s41593-018-0329-4. PubMed PMID: 30742116; PubMed Central PMCID: PMCPMC6508667.

58. Bracko O, Njiru BN, Swallow M, Ali M, Haft-Javaherian M, Schaffer CB. Increasing cerebral blood flow improves cognition into late stages in Alzheimer’s disease mice. J Cereb Blood Flow Metab. 2019:271678×19873658. Epub 2019/09/10. doi: 10.1177/0271678X19873658. PubMed PMID: 31495298.

59. Bailey DM, Marley CJ, Brugniaux JV, Hodson D, New KJ, Ogoh S, et al. Elevated aerobic fitness sustained throughout the adult lifespan is associated with improved cerebral hemodynamics. Stroke. 2013;44(11):3235-8. Epub 2013/08/22. doi: 10.1161/STROKEAHA.113.002589. PubMed PMID: 23963329.

60. Barnes JN, Taylor JL, Kluck BN, Johnson CP, Joyner MJ. Cerebrovascular reactivity is associated with maximal aerobic capacity in healthy older adults. J Appl Physiol (1985). 2013;114(10):1383-7. Epub 2013/03/09. doi: 10.1152/japplphysiol.01258.2012. PubMed PMID: 23471946; PubMed Central PMCID: PMCPMC3656423.

61. Simioni C, Zauli G, Martelli AM, Vitale M, Sacchetti G, Gonelli A, et al. Oxidative stress: role of physical exercise and antioxidant nutraceuticals in adulthood and aging. Oncotarget. 2018;9(24):17181-98. Epub 2018/04/24. doi: 10.18632/oncotarget.24729. PubMed PMID: 29682215; PubMed Central PMCID: PMCPMC5908316.

62. Joyner MJ, Casey DP. Regulation of increased blood flow (hyperemia) to muscles during exercise: a hierarchy of competing physiological needs. Physiol Rev. 2015;95(2):549-601. Epub 2015/04/03. doi: 10.1152/physrev.00035.2013. PubMed PMID: 25834232; PubMed Central PMCID: PMCPMC4551211.

63. Magyar MT, Valikovics A, Czuriga I, Csiba L. Changes of cerebral hemodynamics in hypertensives during physical exercise. J Neuroimaging. 2005;15(1):64-9. Epub 2004/12/03. doi: 10.1177/1051228404269492. PubMed PMID: 15574576.

64. Pologruto TA, Sabatini BL, Svoboda K. ScanImage: flexible software for operating laser scanning microscopes. Biomed Eng Online. 2003;2:13. Epub 2003/06/13. doi: 10.1186/1475-925X-2-13. PubMed PMID: 12801419; PubMed Central PMCID: PMCPMC161784.

65. Santisakultarm TP, Cornelius NR, Nishimura N, Schafer AI, Silver RT, Doerschuk PC, et al. In vivo two-photon excited fluorescence microscopy reveals cardiac- and respiration-dependent pulsatile blood flow in cortical blood vessels in mice. Am J Physiol Heart Circ Physiol. 2012;302(7):H1367-77. Epub 2012/01/24. doi: 10.1152/ajpheart.00417.2011. PubMed PMID: 22268102; PubMed Central PMCID: PMCPMC3330793.

66. Haft-Javaherian M, Fang L, Muse V, Schaffer CB, Nishimura N, Sabuncu MR. Deep convolutional neural networks for segmenting 3D in vivo multiphoton images of vasculature in Alzheimer disease mouse models. PLoS One. 2019;14(3):e0213539. Epub 2019/03/14. doi: 10.1371/journal.pone.0213539. PubMed PMID: 30865678; PubMed Central PMCID: PMCPMC6415838.

67. LeeT.C. KR, L.Chu C.N. Building Skeleton Models via 3-D Medial Surface Axis Thinning Algorithms. CVGIP: Graphical Models and Image Processing. 2002;56(6):Pages 462–78.

68. Manzanares G, Brito-da-Silva G, Gandra PG. Voluntary wheel running: patterns and physiological effects in mice. Braz J Med Biol Res. 2018;52(1):e7830. Epub 2018/12/13. doi: 10.1590/1414-431X20187830. PubMed PMID: 30539969; PubMed Central PMCID: PMCPMC6301263.

69. Beckervordersandforth R, Rolando C. Untangling human neurogenesis to understand and counteract brain disorders. Curr Opin Pharmacol. 2019;50:67-73. Epub 2020/01/07. doi: 10.1016/j.coph.2019.12.002. PubMed PMID: 31901615.

70. Mu Y, Gage FH. Adult hippocampal neurogenesis and its role in Alzheimer’s disease. Mol Neurodegener. 2011;6:85. Epub 2011/12/24. doi: 10.1186/1750-1326-6-85. PubMed PMID: 22192775; PubMed Central PMCID: PMCPMC3261815.

71. Wirths O. Altered neurogenesis in mouse models of Alzheimer disease. Neurogenesis (Austin). 2017;4(1):e1327002. Epub 2018/03/23. doi: 10.1080/23262133.2017.1327002. PubMed PMID: 29564360; PubMed Central PMCID: PMCPMC5856097.

72. Shepherd A, Zhang TD, Zeleznikow-Johnston AM, Hannan AJ, Burrows EL. Transgenic Mouse Models as Tools for Understanding How Increased Cognitive and Physical Stimulation Can Improve Cognition in Alzheimer’s Disease. Brain Plast. 2018;4(1):127-50. Epub 2018/12/20. doi: 10.3233/BPL-180076. PubMed PMID: 30564551; PubMed Central PMCID: PMCPMC6296266.

73. Vecchio LM, Meng Y, Xhima K, Lipsman N, Hamani C, Aubert I. The Neuroprotective Effects of Exercise: Maintaining a Healthy Brain Throughout Aging. Brain Plast. 2018;4(1):17-52. Epub 2018/12/20. doi: 10.3233/BPL-180069. PubMed PMID: 30564545; PubMed Central PMCID: PMCPMC6296262.

74. Ahlskog JE, Geda YE, Graff-Radford NR, Petersen RC. Physical exercise as a preventive or disease-modifying treatment of dementia and brain aging. Mayo Clin Proc. 2011;86(9):876-84. Epub 2011/09/01. doi: 10.4065/mcp.2011.0252. PubMed PMID: 21878600; PubMed Central PMCID: PMCPMC3258000.

75. Buchman AS, Boyle PA, Yu L, Shah RC, Wilson RS, Bennett DA. Total daily physical activity and the risk of AD and cognitive decline in older adults. Neurology. 2012;78(17):1323-9. Epub 2012/04/21. doi: 10.1212/WNL.0b013e3182535d35. PubMed PMID: 22517108; PubMed Central PMCID: PMCPMC3335448.

76. Radde R, Bolmont T, Kaeser SA, Coomaraswamy J, Lindau D, Stoltze L, et al. Abeta42-driven cerebral amyloidosis in transgenic mice reveals early and robust pathology. EMBO Rep. 2006;7(9):940-6. Epub 2006/08/15. doi: 10.1038/sj.embor.7400784. PubMed PMID: 16906128; PubMed Central PMCID: PMCPMC1559665.

77. Serneels L, Van Biervliet J, Craessaerts K, Dejaegere T, Horre K, Van Houtvin T, et al. gamma-Secretase heterogeneity in the Aph1 subunit: relevance for Alzheimer’s disease. Science. 2009;324(5927):639-42. Epub 2009/03/21. doi: 10.1126/science.1171176. PubMed PMID: 19299585; PubMed Central PMCID: PMCPMC2740474.

78. Cui MY, Lin Y, Sheng JY, Zhang X, Cui RJ. Exercise Intervention Associated with Cognitive Improvement in Alzheimer’s Disease. Neural Plast. 2018;2018:9234105. Epub 2018/05/02. doi: 10.1155/2018/9234105. PubMed PMID: 29713339; PubMed Central PMCID: PMCPMC5866875.

79. Winters BD, Forwood SE, Cowell RA, Saksida LM, Bussey TJ. Double dissociation between the effects of peri-postrhinal cortex and hippocampal lesions on tests of object recognition and spatial memory: heterogeneity of function within the temporal lobe. J Neurosci. 2004;24(26):5901-8. Epub 2004/07/02. doi: 10.1523/JNEUROSCI.1346-04.2004. PubMed PMID: 15229237.

80. Assini FL, Duzzioni M, Takahashi RN. Object location memory in mice: pharmacological validation and further evidence of hippocampal CA1 participation. Behav Brain Res. 2009;204(1):206-11. Epub 2009/06/16. doi: 10.1016/j.bbr.2009.06.005. PubMed PMID: 19523494.

81. Barker GR, Warburton EC. When is the hippocampus involved in recognition memory? J Neurosci. 2011;31(29):10721-31. Epub 2011/07/22. doi: 10.1523/JNEUROSCI.6413-10.2011. PubMed PMID: 21775615; PubMed Central PMCID: PMCPMC6622630.

82. Svensson M, Andersson, E., Manouchehrian, O., Yang Y., and Deierborg T. Voluntary running does not reduce neuroinflammation or improve non-cognitive behavior in the 5xFAD mouse model of Alzheimer’s disease. Sci Rep. 2020;10. doi: https://doi.org/10.1038/s41598-020-58309-8.

83. Hoffmann K, Sobol NA, Frederiksen KS, Beyer N, Vogel A, Vestergaard K, et al. Moderate-to-High Intensity Physical Exercise in Patients with Alzheimer’s Disease: A Randomized Controlled Trial. J Alzheimers Dis. 2016;50(2):443-53. Epub 2015/12/20. doi: 10.3233/JAD-150817. PubMed PMID: 26682695.

84. van der Kleij LA, Petersen ET, Siebner HR, Hendrikse J, Frederiksen KS, Sobol NA, et al. The effect of physical exercise on cerebral blood flow in Alzheimer’s disease. Neuroimage Clin. 2018;20:650-4. Epub 2018/09/14. doi: 10.1016/j.nicl.2018.09.003. PubMed PMID: 30211001; PubMed Central PMCID: PMCPMC6129739.

85. Groot C, Hooghiemstra AM, Raijmakers PG, van Berckel BN, Scheltens P, Scherder EJ, et al. The effect of physical activity on cognitive function in patients with dementia: A meta-analysis of randomized control trials. Ageing Res Rev. 2016;25:13-23. Epub 2015/11/27. doi: 10.1016/j.arr.2015.11.005. PubMed PMID: 26607411.

86. Eggermont L, Swaab D, Luiten P, Scherder E. Exercise, cognition and Alzheimer’s disease: more is not necessarily better. Neurosci Biobehav Rev. 2006;30(4):562-75. Epub 2005/12/20. doi: 10.1016/j.neubiorev.2005.10.004. PubMed PMID: 16359729.

87. Alfini AJ, Weiss LR, Nielson KA, Verber MD, Smith JC. Resting Cerebral Blood Flow After Exercise Training in Mild Cognitive Impairment. J Alzheimers Dis. 2019;67(2):671-84. Epub 2019/01/15. doi: 10.3233/JAD-180728. PubMed PMID: 30636734; PubMed Central PMCID: PMCPMC6444938.

88. Sweeney MD, Sagare AP, Zlokovic BV. Blood-brain barrier breakdown in Alzheimer disease and other neurodegenerative disorders. Nat Rev Neurol. 2018;14(3):133-50. Epub 2018/01/30. doi: 10.1038/nrneurol.2017.188. PubMed PMID: 29377008; PubMed Central PMCID: PMCPMC5829048.

89. Iadecola C. The Neurovascular Unit Coming of Age: A Journey through Neurovascular Coupling in Health and Disease. Neuron. 2017;96(1):17-42. Epub 2017/09/29. doi: 10.1016/j.neuron.2017.07.030. PubMed PMID: 28957666; PubMed Central PMCID: PMCPMC5657612.

90. Niwa K, Kazama K, Younkin L, Younkin SG, Carlson GA, Iadecola C. Cerebrovascular autoregulation is profoundly impaired in mice overexpressing amyloid precursor protein. Am J Physiol Heart Circ Physiol. 2002;283(1):H315-23. Epub 2002/06/14. doi: 10.1152/ajpheart.00022.2002. PubMed PMID: 12063304.

