## Supplemental figures 1 and 2 for "Voluntary running does not increase capillary blood flow but promotes neurogenesis and short-term memory in the APP/PS1 mouse model of Alzheimer’s disease"

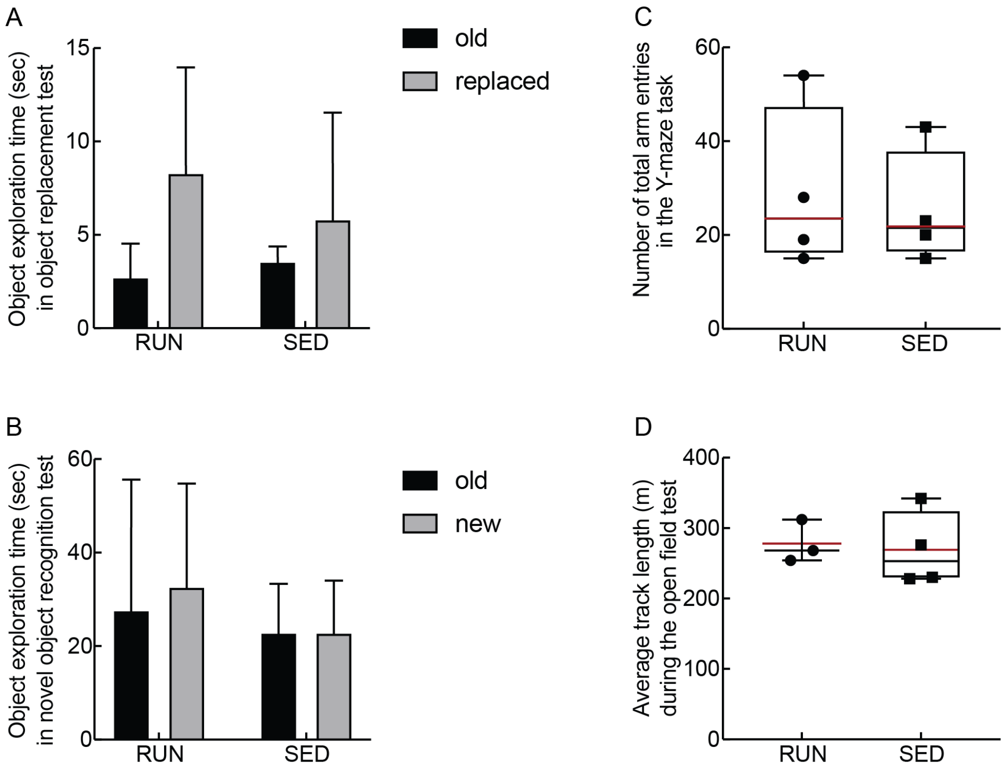


**Fig S1. Running did not affect locomotor activity in behavioral assays.** Both running (RUN) and sedentary (SED) APP/PS1 mice exhibited robust exploration, as assayed by the time spent exploring the old and replaced or new objects, in the **A** object replacement test, and **B** novel object recognition test. Running and sedentary mice also had **C** the same number of arm entries, on average, in the Y-maze task, and **D** the same average total track length in the open field test.

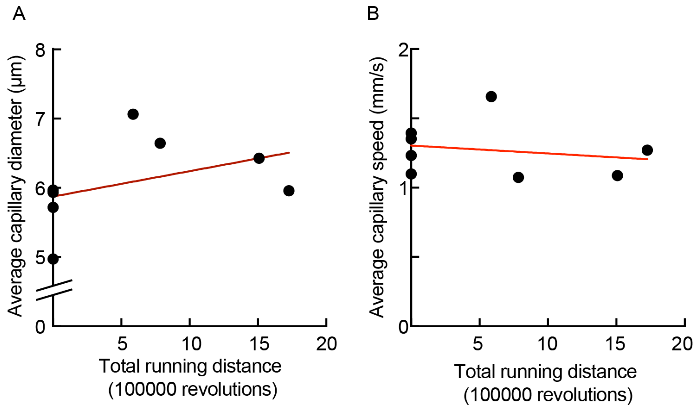


**Fig S2. No correlation was found between the total running distance and the A average capillary diameter or, B average capillary flow speed.**
